# Testicular toxicity following chronic codeine administration is via oxidative DNA damage and up-regulation of NO and caspase 3

**DOI:** 10.1101/796110

**Authors:** RE Akhigbe, A.F Ajayi

## Abstract

**Background:** Codeine, a 3-methylmorphine, and other related opioids have been implicated in androgen suppression, although the associated mechanisms remain unclear.

**Aim:** Therefore, the objective of the current study was to elucidate the in vivo molecular mechanisms underlying codeine-induced androgen suppression.

**Methods:** This study made use of Twenty-one healthy male rabbits, distributed into three groups randomly, control and codeine-treated groups. The control had 1ml of normal saline daily *p.o*. The codeine-treated groups received either 4mg/kg b.w of codeine or 10mg/kg b.w of codeine *p.o*. for six weeks. Reproductive hormonal profile, testicular weight, enzymes, oxidative and inflammatory parameters, histological examination and apoptosis marker were evaluated to examine the effects of codeine use.

**Key findings:** Oral administration of codeine resulted in testicular atrophy and alterations in testicular histomorphology, elevated testicular enzymes, and suppression of circulatory and intra-testicular testosterone. These changes were associated with a marked rise in oxidative markers, including oxidative DNA damage, inflammatory response, and caspase-dependent apoptosis.

**Significance:** In conclusion, chronic codeine use resulted in testicular degeneration and testosterone suppression, which may be attributable to nitric oxide-/oxidativestress-mediated caspase-dependent apoptotic testicular cell death.

## Introduction

Maintenance of fertility is of great concern in the male population, particularly with the rise in the cases of infertility due to male factor. Among the risk factors of male infertility are drug abuse and chronic drug use. Drugs have been shown to impair male fertility via hormonal and non-hormonal mechanisms [1]. They have been reported to suppress the hypothalamic-pituitary-testicular axis resulting in reduced biosynthesis of testosterone [2-5], and consequent decline in spermatogenesis. Some cause direct damage to the sperm cells [6, 7] while others block spermatogenesis [8].

Opioids are among the medications most frequently abused worldwide for self-management of pain and recreational purposes [9-11]. Prolong use of prescription-based opioids can lead to misuse, dependence, and mortality due to their rewarding and reinforcing properties [12], which has made the misuse of pharmacological opioids a growing concern. The prevalence of opioid abuse and misuse among adult opioid users in the United States was 12.8% and 12.5% in 2015 and 2016, respectively [13]. In 2016, about 34.3 million people, accounting for 0.7% of the global population, abused opioids [14].

Chronic use of opioids can lead to androgen deficiency [15-18]. Findings of Rubinstein and Carpenter [15] demonstrated that the highest odds of androgen deficiency are associated with opioids that maintain the most stable serum drug concentrations. Since the biosynthesis of testosterone occurs in a pulsatile manner, it is possible that the nadirs in the concentration of the drug that occurs more between doses of short-acting opioids permit testosterone production that is sufficient to maintain the constant circulating concentration of the androgen [15]. Contrary to this, long-acting opioids suppress testosterone synthesis more completely [15].

Although opioids can cause testicular toxicity, mechanisms associated with opioid-induced testosterone suppression remain unclear. Studies of Azari *et al.* [19] reported that tramadol destroyed testicular germinal layer and atrophy of seminiferous tubules. Their study also revealed that tramadol impaired spermatogenesis and led to necrosis of spermatocytes with nuclear changes such as karyolysis and karyorrhexis. Findings of Nna *et al.* [20] corroborate this report. They observed reduced Leydig cell, testicular germ cell and epididymal spermatozoa density in tramadol-treated rats.

Opioids differ in structure and lipophilicity [12, 15], hence may show variations in their mode of actions. Codeine has been reported to be the most commonly abused opioids globally [21]. Bakare and Isah [22] also reported a similar prevalence among Nigerian abusers. Despite the high prevalence of codeine abuse, there is a dearth of knowledge on the effect of chronic codeine use on testis and testosterone bioavailability. This study sought to demonstrate the possible mechanisms associated with codeine-induced testosterone suppression.

## Methods

### Drugs and chemicals

ELISA kits used for the analysis of reproductive hormones were from Monobind Inc., USA. ELISA kits for analysis of 8-hydroxy-2’-deoxyguanosine (8OHdG), Advanced Glycation endproducts (AGE), and caspase 3 were from Elabscience Biotechnology Co., Ltd, USA. All other reagents not mentioned above were of analytical grade and obtained from Sigma Chemical Co, USA.

### Experimental animals

Twenty-one male rabbits of comparable weight and age (13±2 weeks, 950±50g) housed in standard well-ventilated cages (maximum of 2 animals/cage), in the Animal Holdings of the Department of Physiology, Ladoke Akintola University of Technology, Ogbomoso, Nigeria. The rabbits were provided with standard chow and water *ad libitum* and subjected to the natural photoperiod of 12 h light/dark cycle. All animals were cared for humanely according to the conditions stated in the Guide for the Care and Use of Laboratory Animals’ of the National Academy of Science (NAS), published by the National Institute of Health. The experiment followed the US NAS guidelines and ethical approval obtained from the Ethics Review Committee, Ministry of Health, Oyo State, Nigeria, with Reference Number AD13/479/1396.

### Experimental procedure

Animals were randomly allocated into three groups of seven rabbits each and administered the following:

Group I: control rats received 1ml of 0.9% normal saline.

Group II: low-dose codeine, received 4mg/kg b.w of codeine.

Group III: high-dose codeine, received 10mg/kg b.w of codeine.

All administrations were done orally using oro-pharyngeal cannula daily for six weeks. The 4mg/kg dose was a Human Equivalent Dose, while the 10mg/kg was chosen from dose-response curves to obtain a submaximal peak dose.

Twenty four hours after the last treatment, the over-night-fasted rabbits were weighed and sacrificed via intraperitoneal administration of ketamine (40mg/kg) and xylazine (4mg/kg) [23], and collection of blood samples was via cardiac puncture and centrifuged at 3000rpm for 10 minutes, and the serum separated for hormonal assay. Both testes were excised. Surrounding structures and adipose tissues were trimmed off, and the paired testicular weight of each animal was determined using a sensitive electronic scale. The left testes were kept in sample bottles containing phosphate buffer for further biochemical assay [24, 25]. The testes were homogenized in the buffer, and the homogenate was centrifuged at 10,000rpm for 15 minutes at 4°C to obtain the supernatant for biochemical assay. The removed right testes were fixed in Bouin’s solution for histological processing [26, 27].

### Estimation of testicular enzymes

The testicular activity of alkaline phosphatase (ALP) was determined using a standard kit (Teco Diagnostics, USA). Briefly, testicular tissue was homogenized in phosphate buffer. For each sample, 0.5mL of ALP substrate was dispensed into labeled test tubes and equilibrated to 37°C for 3 minutes. At timed interval, 50µl of each standard, control, and testicular homogenates were added to appropriate test tube. Deionized water was used as sample for reagent blank. The solution was mixed gently and incubated for 10 minutes at 37°C. Following this sequence, 2.5mL of ALP colour developer was added at timed interval and mixed thoroughly. The wavelength of the spectrophotometer was set at 590nm zero with reagent blank (wavelength range: 580-630nm). The absorbance of each sample was read and recorded.

The activity of acid phosphatase (ACP) was determined using a standard kit (Pointe Scientific Inc., USA). Immediately after separation of the supernatant, ACP was stabilized by adding 20µl of Acetate Buffer per 1.0ml of supernatant. The solution was mixed and stored in refrigerator until assay was performed. 1.0 ml of reagent was added to appropriately labeled tube, then 10 µl of L-Tartrate reagent was added and properly mixed. The spectrophotometer was zeroed with water at 405nm, and cuvette temperature set to 37°C. 100 µl of each sample was added, mixed and incubated for 5 minutes. After incubation, the absorbance was read and recorded every minute for five minutes to determine ΔA/minute. Values (U/L) were obtained by multiplying ΔA/minute by the factor.

The activity of lactate dehydrogenase (LDH) was determined using a standard kit (Agappe Diagnostics Ltd., India). 1000µL of working reagent and 10µL of testicular homogenate was mixed and incubated at 37°C for 1 minute. The change in colour absorbance (ΔOD/min) was measured every minute for 3 minutes. The activity of the enzyme was calculated using the formula: LDH-P activity (U/L) = (ΔOD/min) × 16030

The activity of gamma glutamyl transferase (GGT) was determined using a standard kit (Pointe Scientific Inc., USA). The reagents were prepared according to the instruction manual. 1ml of the reagent was pipetted into appropriately labeled tubes: “control”, “sample”, and pre-incubated at 37°C for 5 minutes. The spectrophotometer was zeroed with water at 405nm. 100 µl of the homogenate was added, mixed and returned to a thermo cuvette. After 60 seconds, the absorbance was read and recorded every minute for 2 minutes. The mean absorbance difference per minute (Δ Abs./min.) was determined and multiplied by 1158 to obtain results in U/L.

### Estimation of testicular total protein and glucose

Testicular total protein concentration was measured using the biuret method [28], which depends on the presence of peptide bonds in all proteins. These peptide bonds react with Cu^2+^ ions in alkaline solutions to form a coloured product. The absorbance of which is measured spectrophotometrically at 540nm.

Testicular glucose concentration was determined using standard kit (Agappe Diagnostics Ltd., India). 1000µL of the working reagent and 10µL of the testicular homogenate were mixed and incubated for 10 minutes at 37°C. The absorbance of standard and sample were read against reagent blank, and the concentration of glucose determined by the formula:

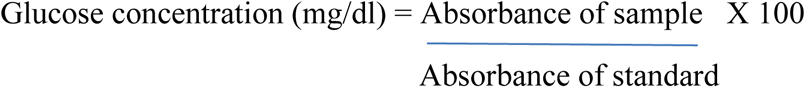

### Estimation of serum and testicular levels of reproductive hormones

Serum and testicular testosterone were determined using ELISA kit (Monobind Inc. USA; product number: 4806-300A), serum oestrogen was determined using ELISA kit (Monobind Inc. USA; product number: 4925-300A), serum FSH was assayed using ELISA kit (Monobind Inc. USA; product number: 506-300A), and serum LH was assayed using ELISA kit (Monobind Inc. USA; product number: 625-300A). The hormones were assayed as previously documented [25, 26]. Briefly, for serum testosterone assay, 10µL of standards, control and serum samples were dispensed into their respective wells. 100 µL Testosterone-HRP conjugate was added to each well. Substrate blank was dispensed into well A1. The wells were then covered with foil. Then incubation was carried out for 1 hour at room temperature, the foil was removed and well contents aspirated. Then 300 µL diluted wash solution was used three times to wash the wells. The soak time between each wash cycle was more than 5 seconds. The remaining fluid was carefully removed by tapping the strips on tissue paper. 100 µL of TMB substrate solution was added into all wells. The wells were incubated for 15 minutes at room temperature in the dark. 100 µL stop solution was dispensed into all wells in the same order and the same rate as for the substrate. The absorbance of the specimen was read at 450nm within 20 minutes after addition of the stop solution.

Testicular testosterone measurement was determined similarly; however, the testes supernatant obtained from testicular homogenate was used instead of the serum.

Similarly, determination of serum oestrogen was by dispensing 25µL of standards, control and serum samples into their respective wells. 50µL of E2 Biotin reagent was added to all wells and incubated for 30 minutes at 37°C after which 50µL of enzyme conjugate was added to each well. Well A1 was left for substrate blank. The wells were covered with foil. Incubated of the wells was for 90 minutes at room temperature. The foil was removed and well contents aspirated. 300 µL diluted wash solution was used to wash the wells three times. The soak time between each wash cycle was more than 5 seconds. The remaining fluid was carefully removed by tapping the strips on tissue paper. 100µL of TMB substrate was added into all wells. Incubation of the wells was carried out in the dark at 37°C for 30 minutes. 50 µL of stop solution was then dispensed into all wells in the same order and time interval as the substrate. The absorbance of the specimen was read at 450nm within 20 minutes after the addition of the stop solution.

For FSH assay, 50µL of standards, control and serum were dispensed into their respective wells. 100 µL of the conjugate was added to each well. Substrate blank was dispensed into well A1. The wells were covered with foil. Incubation of the wells was carried out for 1 hour at 37°C. After 1 hour the foil was removed and well contents aspirated, wells washed three times with 300 µL of diluted wash solution. The soak time between each wash cycle was more than 5 seconds. The remaining fluid was carefully removed by tapping the strips on tissue paper. 100 µL of TMB substrate was added into all wells. The wells were incubated for 15 minutes at room temperature in the dark. 50µL of stop solution was dispensed into all wells in the same order and rate as for the substrate. The absorbance of the specimen was read at 450nm.

For LH assay, 50µL of standards, control and serum samples were dispensed into their respective wells in duplicates. 100 µL conjugate was added to each well. Substrate blank was dispensed into well A1. The wells were left to incubate for 1 hour at room temperature. Contents of wells were aspirated after 1 hour, and the wells were washed three times using 300 µL diluted wash solution. The remaining fluid was carefully removed by tapping the strips on tissue paper. 100µL of TMB substrate solution added into all wells. The wells were incubated for 15 minutes at room temperature in the dark. 100µL of stop solution was dispensed into all wells in the same order and rate as for the substrate. The absorbance of the specimen was done at 450nm.

The concentrations of the hormones were obtained from the absorbance values using standard curves obtained.

### Estimation of testicular oxidative markers, anti-oxidants, 8OHdG and caspase 3

Lipid peroxidation concentration determination is by measuring the thiobarbituric acid reactive substances (TBARS) produced during lipid peroxidation as previously documented [29]. This method is dependent on the reaction between 2-thiobarbituric acid (TBA) and malondialdehyde, an end product of lipid peroxidation. On heating in acidic pH, a pink chromogen complex ([TBS] 2-malondialdehyde adduct) is formed and measured by its absorbance at 532 nm. Briefly, 200µL of the sample was deproteinised with 500µL of Trichloroacetic acid (TCA) and centrifuged at 3000rpm for 10minutes. 1mL of 0.75% TBA was added to 0.1mL of the supernatant and boiled in a water bath for 20minutes at 100°C and cooled with ice water. The absorbance of the sample/standard was read at 532nm with a spectrophotometer against the blank. The concentration of TBARS generated was extrapolated from the standard curve.

AGE was determined using a standard ELISA kit (Elabscience Biotechnology Co., Ltd, USA; product number: E-EL-010296T) following the manufacturer’s manual. The standard working solution was added to the first two columns: Each concentration of the solution was added in duplicate, to one well each, side by side (50 uL for each well). Samples were added to the other wells (50 uL for each well). Immediately, 50μL of Biotinylated Detection Ab working solution was added to each well. The plate was then covered with the provided sealer in the kit and incubated for 45 minutes at 37°C. Care was made to ensure that the solutions were added to the bottom of the micro ELISA plate well. Also, avoidance of touching the inside wall and causing foaming as much as possible was ensured. The solution aspirated from each well, and 350 uL of wash buffer added to each well. These were soaked for 1-2 minutes, and the solution was aspirated from each well, then patted dry against clean absorbent paper. The wash step was repeated three times. 100 μL of HRP Conjugate working solution was added to each well and covered with the plate sealer then incubated for 30 minutes at 37°C. The solution from each well was aspirated, and the wash process was repeated five times. Then 90 μL of Substrate Reagent was added to each well and covered with a new plate sealer then incubated for another 15 minutes at 37°C. The plate was protected from light. After this, 50 μL of stop solution was added to each well. The value of the light density of each well was determined with a microplate reader set to 450 nm.

8OHdG and caspase three levels were determined as described for AGE using ELISA kits (Elabscience Biotechnology Co., Ltd, USA) with product numbers E-EL-0028 and E-EL-RB0656, respectively.

Hydrogen peroxide (H_2_O_2_) generation was determined by the method described by Nouroozzadeh *et al.* [30]. The method uses a colour reagent that contains xylenol orange dye in an acidic solution with sorbitol and ferrous ammonium sulphate that reacts to produce a purple colour in proportion to the concentration of H_2_O_2_. The assay mixture was thoroughly vortex-mixed until it foamed. A pale pink colour complex was generated after incubation for 30 minutes at room temperature. The absorbance was read against blank (distilled water) at 560nm wavelength. The concentration of hydrogen peroxide generated was extrapolated from the standard curve.

Myeloperoxidase activity was determined as previously documented by Desser *et al*. [31]. Myeloperoxidase catalyses the oxidation of guaiacol to oxidised guaiacol in the presence of hydrogen peroxide. Guaiacol in its oxidised form has a brown colour which is measured photometrically at 470nm wavelength. The concentration of oxidised guaiacol produced in the reaction is proportional to the colour intensity produced.

Superoxide dismutase (SOD) activity was determined as previously reported [32, 33]. Briefly, 1mL of the sample was diluted in 9mL of distilled water to make a 1 in 10 dilutions. An aliquot of 0.2mL of the diluted sample was added to 2.5mL of 0.05M carbonate buffer (pH 10.2) for spectrophotometer equilibration and the reaction started by the addition of 0.3mL of freshly prepared 0.3mM adrenaline to the mixture which was quickly mixed by inversion. The reference cuvette contained 2.5mL buffer, 0.3mL of the substrate (adrenaline) and 0.2mL of water. The increase in absorbance was monitored every 30 seconds for 150 seconds 480nm.

Testicular catalase activity was determined as previously documented [34] with some modifications. Briefly, 1ml of the supernatant of the testicular homogenate was mixed with 19ml of diluted water to give a 1:29 dilution of the sample. The assayed mixture contained 4ml of H_2_O_2_ solution (800µmoles) and 5ml of phosphate buffer in a 10ml flat bottom flask. 1ml of properly diluted enzyme preparation was mixed with the reaction mixture by a gentle swirling motion at 37°C room temperature. 1ml of the reaction mixture was withdrawn and blown into 2ml dichromate/acetic acid reagent at 60 seconds intervals. The H_2_O_2_ contents of the withdrawn sample were determined. Catalase levels in the sample are determined by comparing absorbance at 653 nm to that of a certified catalase standard.

The method of Beutler *et al*. [35] was used in estimating the level of reduced glutathione (GSH). An aliquot of the sample was deproteinized by the addition of an equal volume of 4% sulfosalicylic acid, which was centrifuged at 4,000rpm for 5 minutes. After that, 0.5ml of the supernatant was added to 4.5ml of Ellman’s reagent. A blank was prepared with 0.5ml of the diluted precipitating agent and 4.5ml of Ellman’s reagent. Reduced glutathione, GSH, is equal to the absorbance at 412nm.

Testicular glutathione peroxidase (GPx) activity was determined according to the method of Rotruck *et al*. [36] with some modifications. Briefly, the reaction mixture containing the sample was incubated at 37°C for 3 minutes after which 0.5mL of 10% trichloroacetic acid (TCA) was added and after that centrifuged at 3000rpm for 5 minutes. To 1mL of each of the supernatants, 2mL of phosphate buffer and 1mL of 5’-5’-dithiobis-(2-dinitrobenzoic acid (DTNB) solution was added, and the absorbance was read at 412nm against a blank. Glutathione peroxidase activity was observed by plotting the standard curve, and then the concentration of the remaining GSH extrapolated from the curve.

Testicular Glutathione-S-transferase activity was determined using the method described by Habig *et al*. [37]. The method is based on the relatively high activity of glutathione-S-transferase with 1-chloro-2,4,-dinitrobenzene as the second substrate. The prepared medium of the assay and the reaction was allowed to run for 60 seconds, then the absorbance was read against the blank at 340nm. The temperature was put at approximately 37°C.

Testicular nitric oxide was determined using Griess Reaction as previously documented [38]. 100µL of Griess reagent, 300µL of nitrate-containing testicular homogenate, and 2.6ml of deionized water were mixed in a spectrophotometer cuvette and incubated for 30 minutes at room temperature. The blank was prepared by mixing 100µL of Griess reagent and 2.9ml of deionised water. The absorbance of the nitrate-containing sample was measured at 548nm relative to the reference sample.

### Histology

Sections of the fixed tissues were obtained from paraffin blocks and stained with Haematoxylin and Eosin (H&E) [39]. Stained sections of the testis were observed under a digital light microscope at various magnifications and photographs were taken.

### Statistical analysis

Data were expressed as mean ± SD. The data comparisons made use of one-way analysis of variance (ANOVA) followed by Tukey’s post hoc test for pairwise comparison. Pearson’s bivariate correlation was done to assess the correlation between selected indices of testicular functions and 8-OHdG, nitric oxide and caspase 3. Statistical Package for Social Sciences (SPSS, version 16) was used for the analysis. Statistical significance was set at *p* < 0.05.

## Results

### Effect of chronic codeine use on body weight, testicular weight and relative testicular weight

Table 1 represents the body and testicular weights of the treated and control groups. Oral administration of codeine for six weeks did not statistically affect the body weight gain. However, oral administration of 4mg/kg and 10mg/kg bodyweight of codeine showed 50% and 68% decline in paired testicular weight, respectively. The data on paired testicular weight presented in Table 1 showed a dose-dependent decrease in the paired testicular weight of the treated rabbits. The relative testicular weight determined as the percentage of the ratio between the paired testicular weight and body weight also showed significant differences across the groups. The decline in the relative testicular weight for codeine-treated rabbits at 4mg/kg and 10mg/kg body weight was 50% and 200% respectively below the control rabbits.

**Table 1:** Effect of chronic codeine use on body weight, testicular weight and relative testicular weight.

| Group | Initial BW (g) | Final BW (g) | Difference (g) | Difference (%) | PTW (g) | PTW/BWX 100 |
| --- | --- | --- | --- | --- | --- | --- |
| **Control** | 872.00±69.63<sup>a</sup> | 1379.1±54.51<sup>a</sup> | 507.05±32.93<sup>a</sup> | 58.6257±7.57<sup>a</sup> | 2.84±0.19<sup>a</sup> | 0.21±0.01<sup>a</sup> |
| **Low-dose codeine** | 866.62±60.21<sup>a</sup> | 1353.8±71.95<sup>a</sup> | 487.18±26.15<sup>a</sup> | 56.3971±4.24<sup>a</sup> | 1.40±0.55<sup>b</sup> | 0.11±0.04<sup>b</sup> |
| **High-dose codeine** | 818.91±46.69<sup>a</sup> | 1305.2±64.86<sup>a</sup> | 486.32±26.69<sup>a</sup> | 59.4614±3.22<sup>a</sup> | 0.91±0.38<sup>b</sup> | 0.07±0.03<sup>b</sup> |

Values represent mean±SD for 7 replicates in each group. BW, body weight of rats; PTW, paired testicular weight. Parameters carrying different superscripts are significantly different at $p < 0.05$ .

### Effect of chronic codeine use on testicular enzymes

Figure 1 shows the activities of testicular enzymes (alkaline phosphatase (ALP), acid phosphatase (ACP), lactate dehydrogenase (LDH) and γ-glutamyl-transferase (GGT)) in control and experimental rabbits. The activities of these enzymes were observed to be significantly raised in the treated rabbits when compared with the control rabbits (*p* < 0.05). Oral administration of codeine for six weeks at 4mg/kg body weight led to a 29% rise in ALP, while at 10mg/kg body weight caused a further rise by 6.5%. Similarly, oral codeine administration at 4mg/kg body weight led to a rise in ACP, LDH and GGT by 19%, 13.7%, and 11.8% respectively. At 10mg/kg body weight, codeine administration further increased the activities of LDH, and GGT by 4.4% and 12% respectively, and decreased ACP activity by 0.2% when compared to rabbits who had 4mg/kg body weight of codeine. Although the difference observed in the treated groups were statistically similar, there was a significant difference in GGT activities of both treated groups.

**Figure 1:**
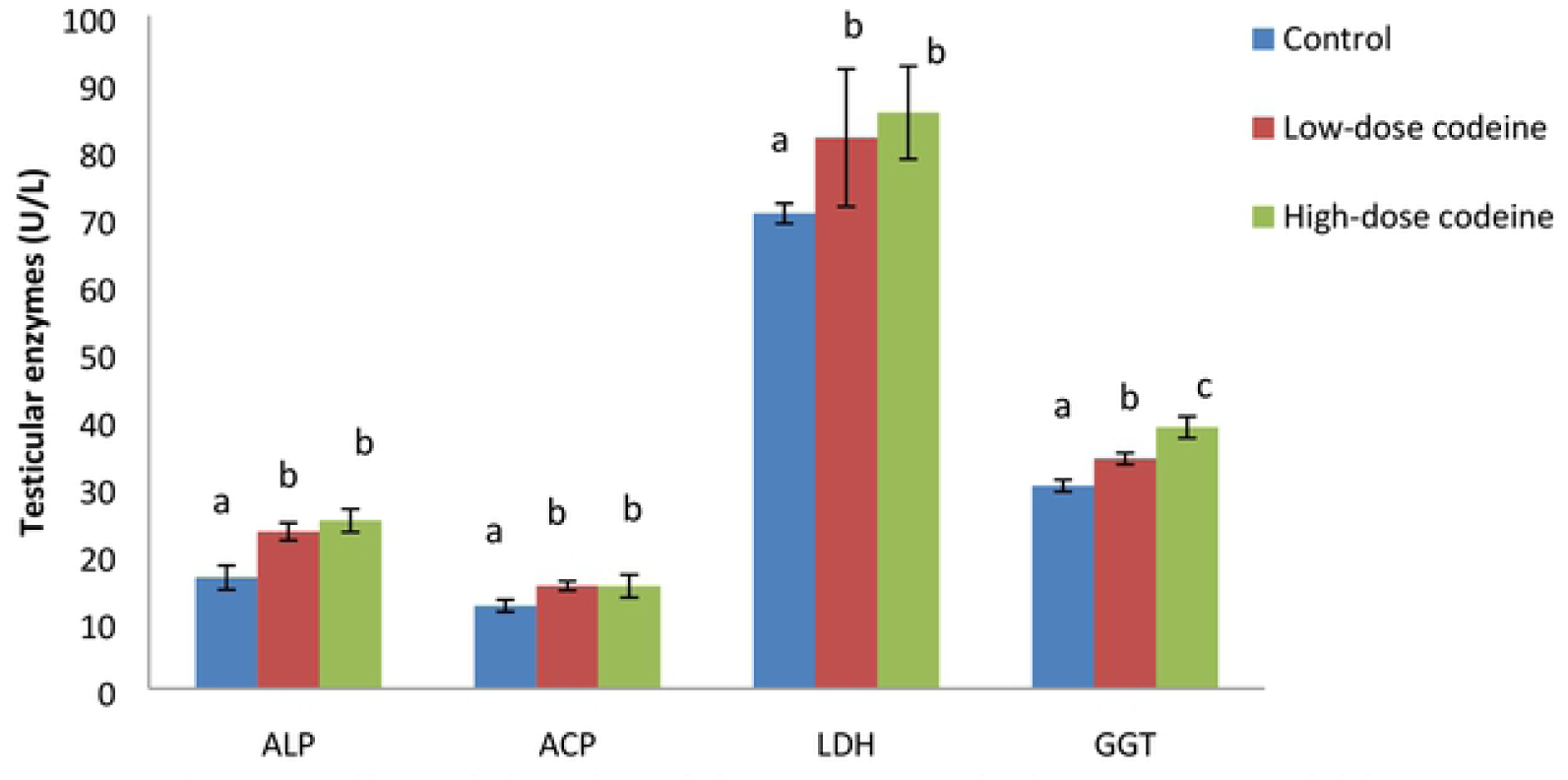
Effect of chronic codeine use on testicular enzymes activities. Bars of the same parameter carrying different alphabets (a,b,c) are statistically different

### Effect of chronic codeine use on testicular total protein and glucose

Figure 2 shows testicular levels of total protein and glucose. Although oral administration of codeine at four and 10mg/kg body weight led to a rise in total testicular protein by 3.8% and 7.5% respectively, and a similar increase in testicular glucose by 12.9% and 18.6% respectively when compared with the control; these changes were marginal across all groups.

**Figure 2:**
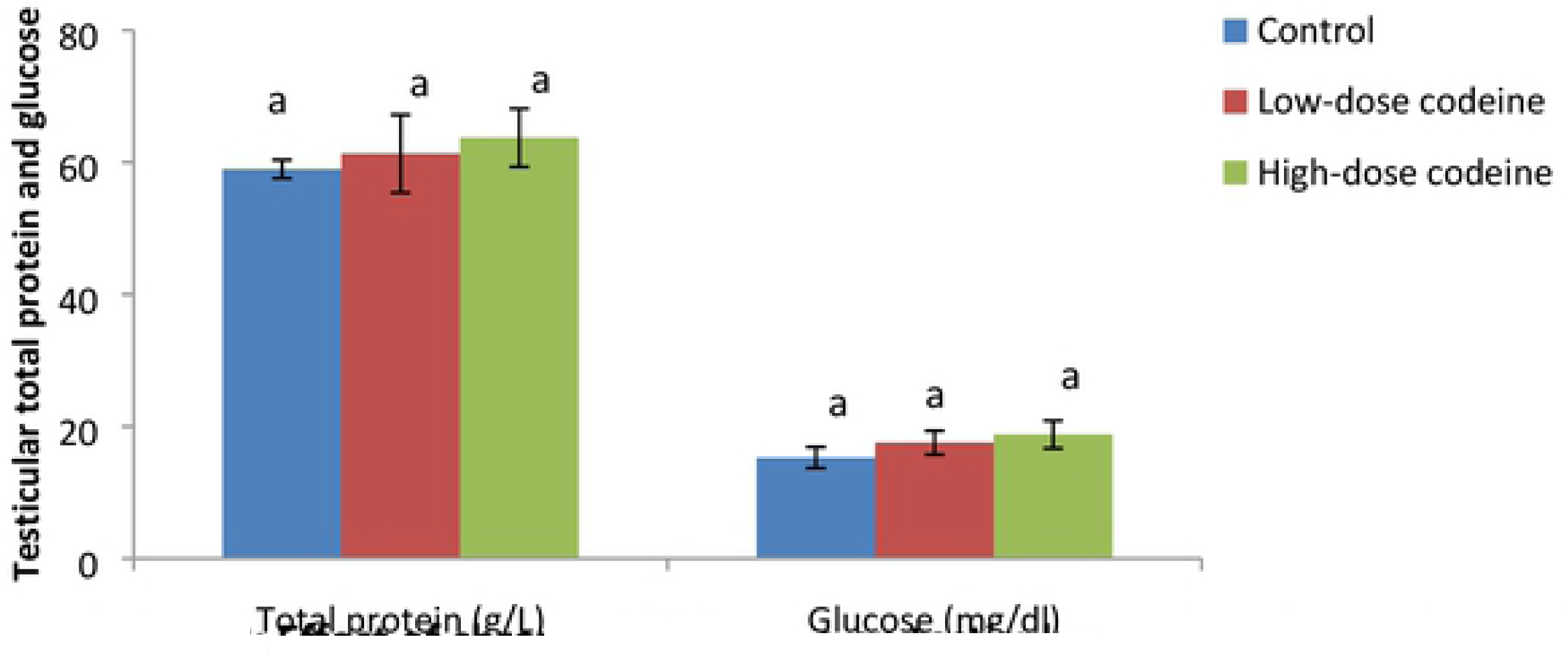
Effectorcnfobic codeine use onTeslFculartotal protein and glucose. Bars of the same parameter carrying different alphabets (a,b,c) are statistically different

### Effect of chronic codeine use on serum and testicular male reproductive hormones

Figure 3 shows the influence of chronic oral administration of codeine on serum concentrations of testosterone, LH, and FSH, and testicular level of testosterone in rabbits. Circulatory concentrations of LH and FSH in codeine-treated animals significantly increased when compared with those of the control animals. However, oral administration of codeine at four and 10mg/kg body weight significantly suppressed circulatory testosterone levels by 24% and 34% respectively when compared with the control. Similarly, codeine treatment at four and 10mg/kg body weight led to a significant dose-dependent decline in the testicular concentration of testosterone by 24.5% and 34.7% when compared with the control.

**Figure 3:**
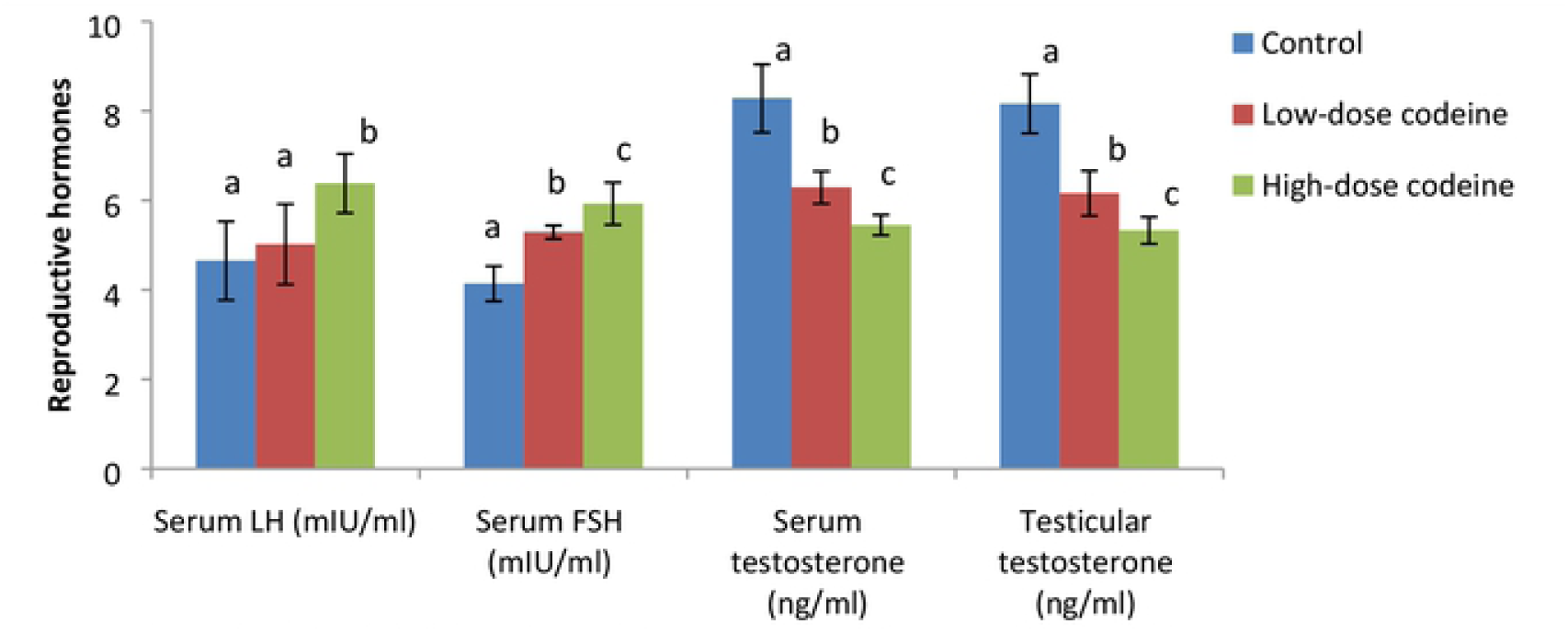
Effect of chronic codeine use on male reproductive hormones. Bars of the same parameter carrying different alphabets (a,b,c) are statistically different

Figure 4 shows the values of hormonal fertility indices. The results showed that the values of serum testosterone/luteinizing hormone (T/LH) and testosterone/oestrogen (T/E_2_) were significantly reduced by codeine exposure. Low dose codeine at 4mg/kg body weight caused 29% and 12% decline in T/LH and T/E_2,_ respectively when compared with the control. Administration of high dose codeine at 10mg/kg body weight caused further decline in these indices by 35% and 2.5% respectively.

**Figure 4:**
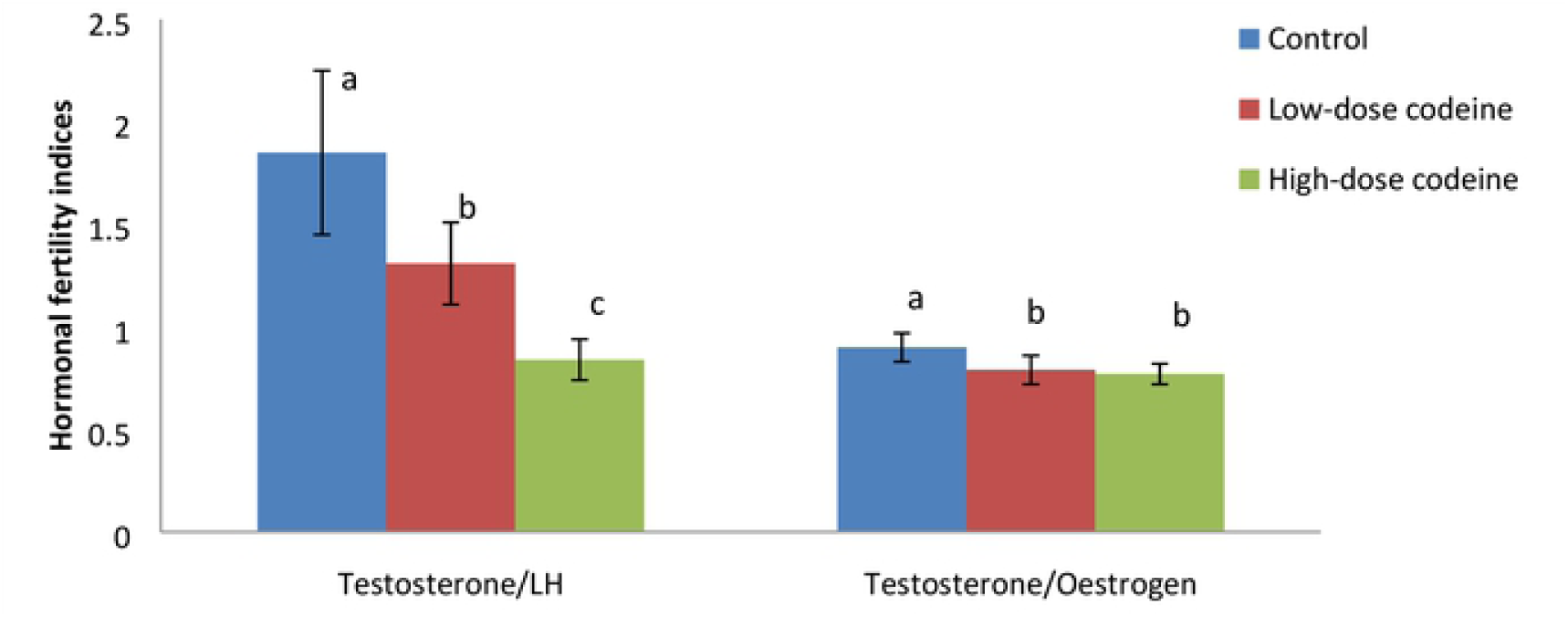
Effect of chronic codeine use on hormonal fertility indices. Bars of the same parameter carrying different alphabets (a,b,c) are statistically different

### Effect of chronic codeine use on testicular oxidative markers and enzymatic anti-oxidants

Oxidative damage induced by oral administration of codeine in testicular tissue resulted in significant dose-dependent rise in the levels of lipid peroxidation index (malondialdehyde, MDA), oxidative protein denaturation index (AGE), hydrogen peroxide and myeloperoxidase (Figure 5). On the other hand, codeine treatment significantly lowered the levels of SOD, catalase, GSH, GPx and GST. These reductions were also dose-dependent (Figure 6).

**Figure 5:**
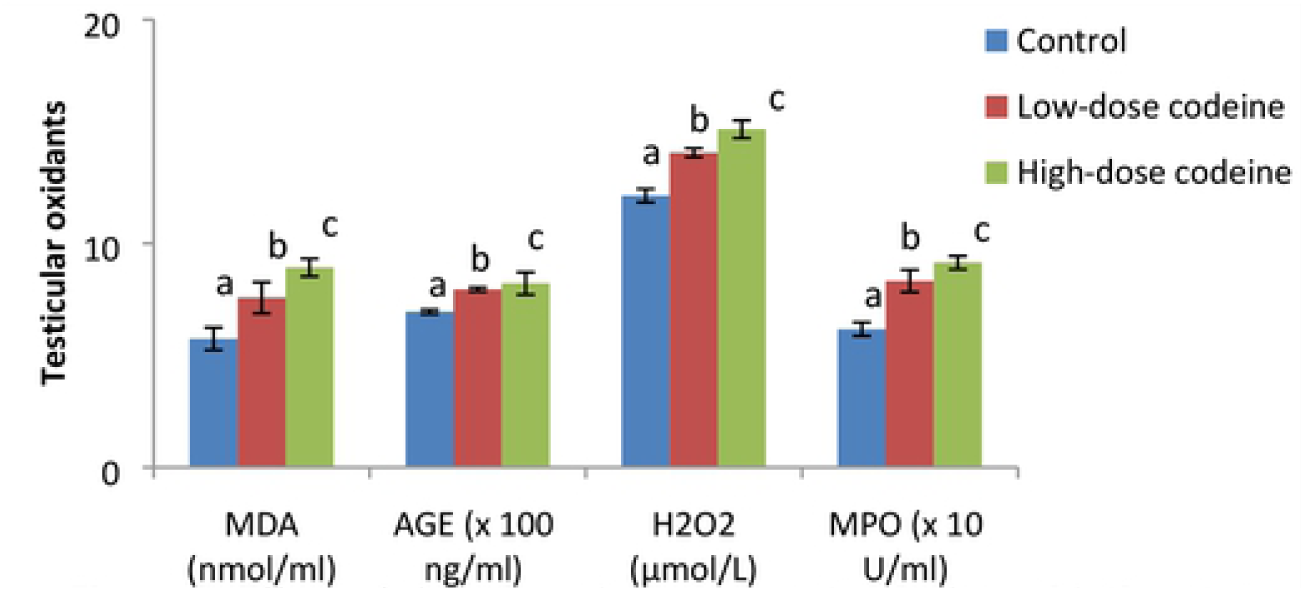
Effect of chronic codeine use on testicular oxidative stress markers. Bars of the same parameter carrying different alphabets (a,b,c) are statistically different

**Figure 6:**
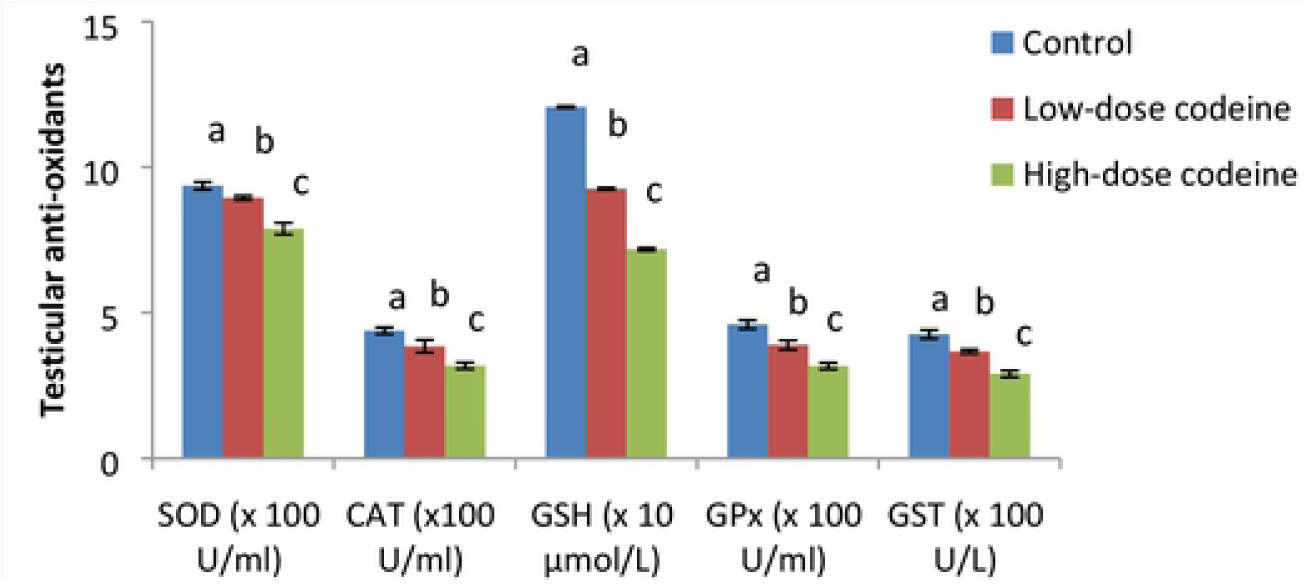
Effect of chronic codeine use on testicular enzymatic anti-oxidants. Bars of the same parameter carrying different alphabets (a,b,c) are statistically different

### Effect of chronic codeine use on testicular DNA damage, nitric oxide and apoptosis

Compared with the control group, testicular oxidative DNA damage significantly increased in the codeine-treated groups (Figure 7). Low-dose codeine at 4mg/kg body weight led to 27% increase in testicular oxidative DNA damage, while high-dose codeine at 10mg/kg body weight led to 41% increase in testicular oxidative DNA damage when compared with the control. Also, a significant increase in the testicular level of nitric oxide was observed in codeine-treated rabbits (Figure 8). The rise in nitric oxide seen was dose-dependent. Oral administration of codeine for six weeks at four and 10mg/kg body weight led to 35.6% and 55.7% increase in testicular nitric oxide level.

**Figure 7:**
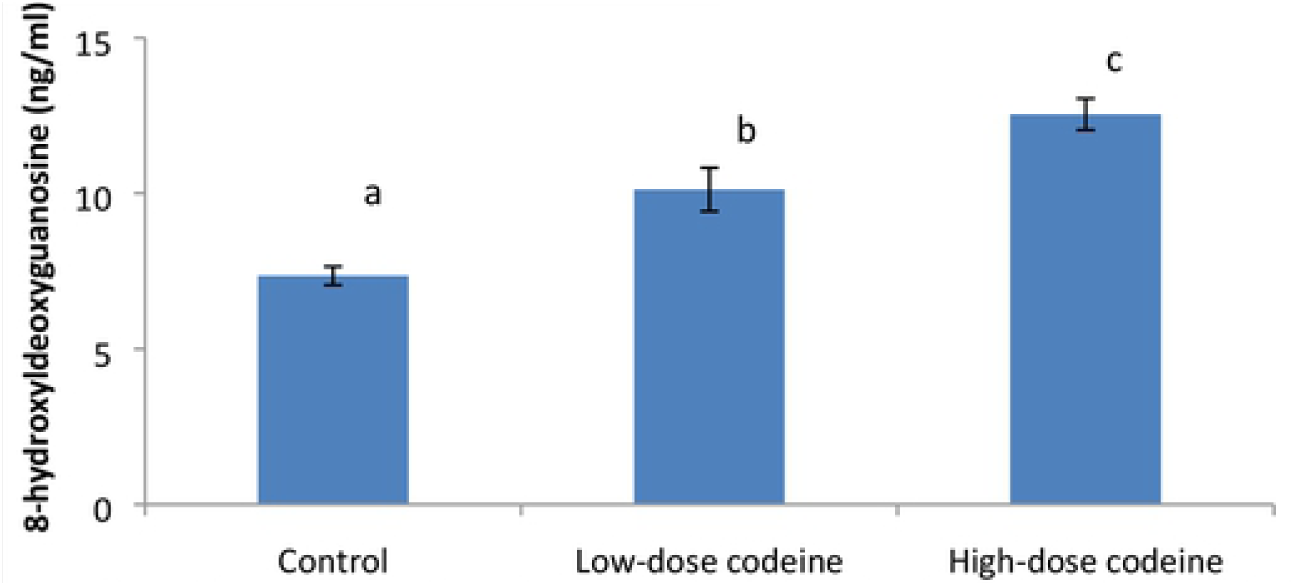
Effect of chronic codeine use on testicular oxidative DNA damage. Bars of the same parameter carrying different alphabets (a,b,c) are statistically different

**Figure 8:**
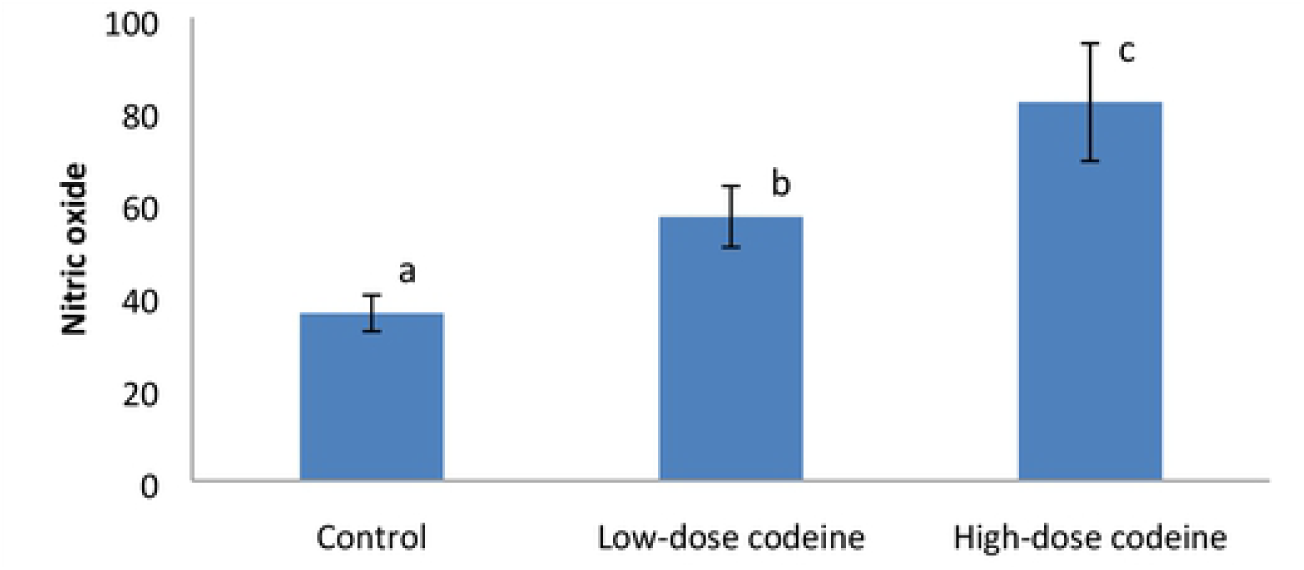
Effect of chronic codeine use on testicular nitric oxide. Bars of the same parameter carrying different alphabets (a,b,c) are statistically different

The assessment of the role of apoptosis was by measuring the concentration of caspase three. Codeine treatment was observed to cause caspase-mediated apoptosis in a dose-dependent manner (Figure 9). At four and 10mg/kg body weight, oral administration of codeine significantly increased the testicular level of caspase three by 66.8% and 81% respectively when compared with the control.

**Figure 9:**
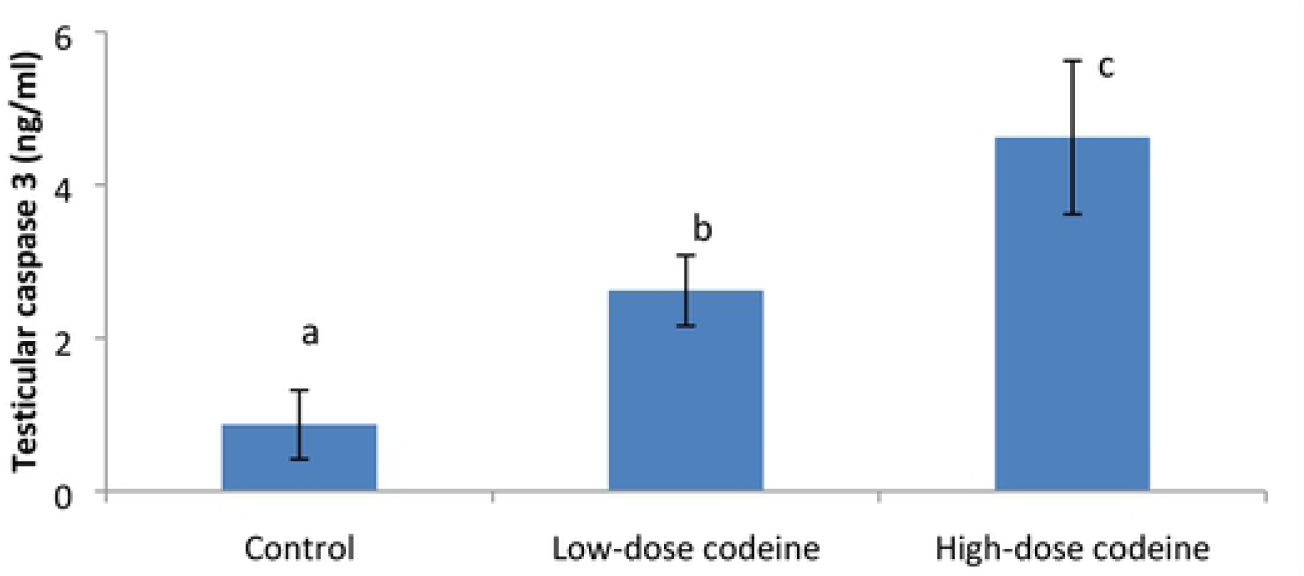
Effect of chronic codeine use on testicular caspase 3. Bars of the same parameter carrying different alphabets (a,b,c) are statistically different

### Correlation study

A negative correlation was observed between testicular weight, serum and intra-testicular testosterone, and hormonal fertility indices, and 8-OHdH, nitric oxide and caspase 3 activity (Table 2).

**Table 2:** Comparison of some fertility indices with markers of oxidative DNA damage, infla mmation, and apoptosis relative t esticular weight.

|  | 8-OHdG | Nitric oxide | Caspase 3 |
| --- | --- | --- | --- |
| **Paired testicular weight** | -0.836 | -0.840 | -0.749 |
| **Relative testicular weight** | -0.823 | -0.824 | -0.713 |
| **Testicular testosterone** | -0.864 | -0.825 | -0.832 |
| **Serum testosterone** | -0.894 | -0.828 | -0.824 |
| **Testosterone/LH** | -0.803 | -0.781 | -0.731 |
| **Testosterone/Oestrogen** | -0.624 | -0.564 | -0.498 |

P < 0.01
Values represent mean±SD for 7 replicates in each group. BW, body weight of rats; PTW, paired testicular weight. Parameters carrying different superscripts are significantly different at p < 0.05.

### Histological findings

Findings of histopathological examination supported the present biochemical findings. Photomicrographs (Figure 10) show control rabbits with normal seminiferous tubules containing healthy germ cells, including spermatogonia cells and Sertoli cells, and normal maturation stages with the presence of spermatozoa within their lumen. The interstitial spaces show healthy Leydig cells. Oral administration of codeine at four and 10mg/kg body weight caused alteration of testicular histoarchitecture. Codeine administration resulted in thickened propria indicative of cessation of spermatogenesis and vascular congestion and associated vacuolation, sloughed germ cells and maturation arrest.

**Figure 10:**
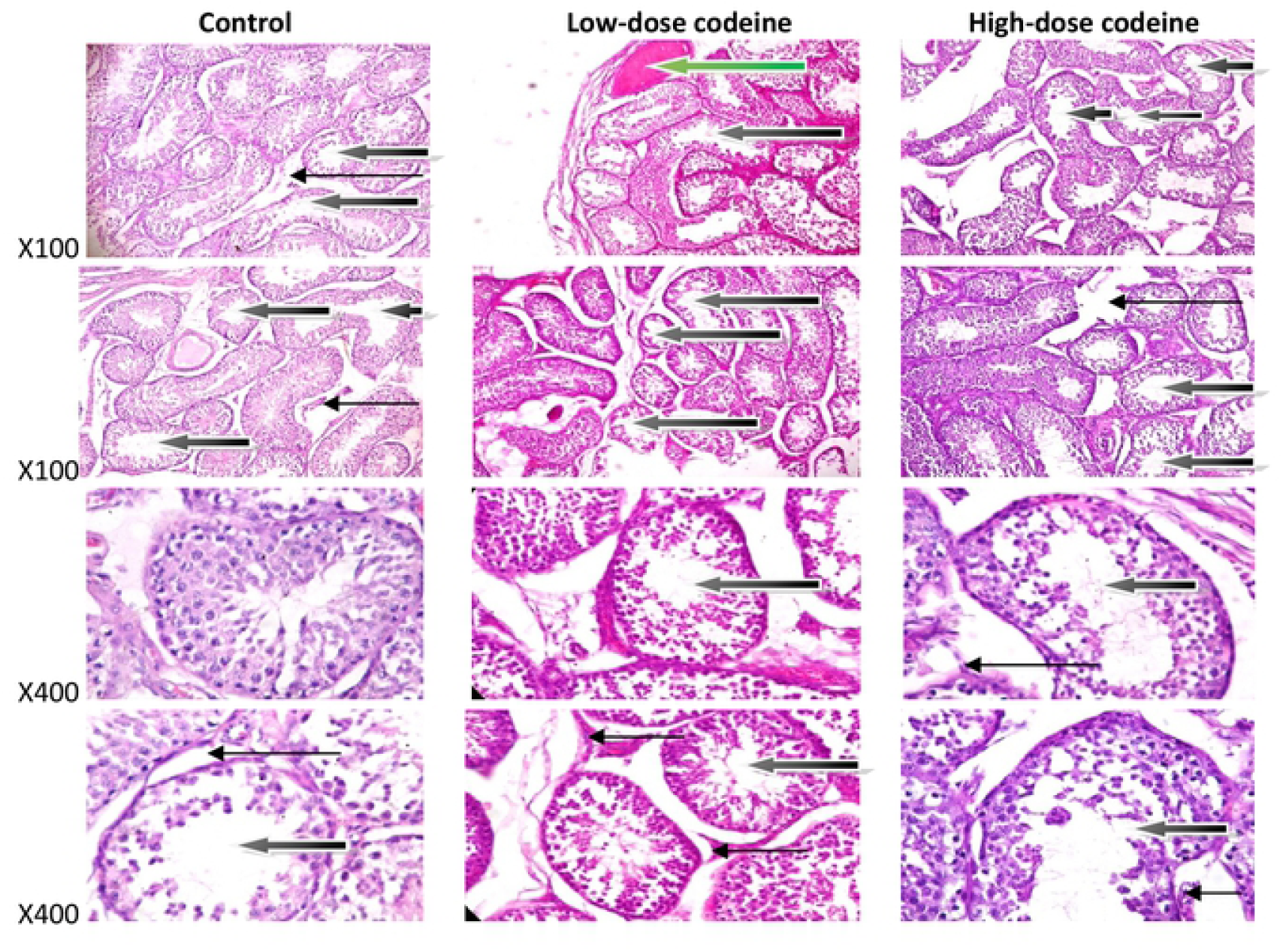
Photomicrographs of testicular sections stained by Haematoxylin and Eosin. The control shows some norm al seminiferous tubules containing normal germ cells including spermatogonia cells and sertoli cells, and normal maturation stages with presence of spermatozoa within their lumen(white arrow). There very few tubules show maturation arrest at secondary stage of maturation (black arrow).The interstit ial spaces show normal leydig cells (slender arrow). The rabbits on low-dose codeine show very poor architecture. The seminiferous tubules show double celllayer or thickened propria indicative of cessation of spermatogenesis. They exhibit vacuolation, sloughed germ cells and maturation arrest and show giant cells (black arrow). There is evidence of vascular congestion (green arrow).The interstitia l spaces appear normal (slender arrow). The animals on high-dose codeine show very poor architecture. The seminiferous tubul es show double cell layer or thickened propria indicative of cessation of sperm atogenesis. They exhibit vacuolation, sloughed germ cells and maturation arrest and show giant cells (black arro w). The intersti tial spaces appear normal (slender arr ow).

## Discussion

The abuse of codeine, a 3-methylmorphine, remains a global health threat. This drug molecule has been reported to suppress testosterone production. However, this study is the first to demonstrate the toxic effect of codeine on the testis and testicular function via oxidative- and inflammation-mediated caspase-dependent apoptosis. Ranging from a reduction in testicular weight, impaired testicular functions, spermatogenesis cessation, reduced hormonal fertility indices, steroidogenesis, as well as induction of testicular oxidative damage, inflammation, and caspase-mediated apoptosis observed in our study, the need to address the menace of opioid abuse would go a long way in tackling the challenges posed by drug-induced infertility.

The reduction in testicular weight observed in this study is a pointer to codeine-induced alteration in metabolic process that could be associated with excessive tissue break down [40, 41]. Studies have reported the possibility of assessing organ toxicity by the organ weight [42, 43]. Though there was a similar body weight change in the treated and control animals, the decrease in testicular weight, a valuable index of reproductive toxicity, seen in the codeine-treated rabbits indicates that the drug molecule has a toxic effect on the testis and consequent depression of testicular metabolism and function. The reduction in testicular weight following codeine use explains the associated reduction of testosterone bioavailability and disrupted spermatogenesis observed in this study. The decline in the circulatory concentrations of testosterone seen in codeine treatment aligns with previous reports [15-18]. The decline in serum and testicular levels of androgen and cessation of spermatogenesis observed in codeine-treated animals could be partly due to codeine-induced loss of testicular tissues, and impairment of testicular metabolism and function.

Codeine administration induced oxidative stress in the rabbit testis, evident by enhanced lipid peroxidation, oxidative protein denaturation and compromised enzymatic anti-oxidant defence system revealed by decreased glutathione systems with a concomitant reduction in superoxide dismutase and catalase. These findings agree with the report of Nna and Osim [44] that demonstrated the oxidative effects of tramadol on rat testis. Oxidative stress is as a result of an imbalance between reactive oxygen species (ROS) generation and intracellular ROS scavenging capacity. Although the increase in testicular oxidative stress has been reported following use of opioid-like tramadol [44], no study has reported whether or not codeine shares the same activity. The testes are quite prone to ROS-induced injury and peroxidative damage consequent to their high concentrations of polyunsaturated fatty acids and low antioxidant capacity [45].

Reduced glutathione (GSH) scavenges ROS by forming oxidized glutathione (GSSG) and other disulfides. Increased oxidative stress promotes the formation and efflux of GSSG, and glutathione reductase (GR) mediates the reduction of GSSG to GSH [46]. This reaction requires the reduced form of nicotinamide adenine dinucleotide phosphate (NADPH), which is provided by glucose-6-phosphate dehydrogenase (G6PD) of the pentose phosphate pathway [46]. Codeine-induced decline in the activity of GSH is indisputably likely consequent to reduced G6PD activity and the resultant decrease in the availability of NADPH. GPx is another constituent of the GSH redox cycle. It is the primary enzyme responsible for hydrogen peroxide (H_2_O_2_) elimination in the testis [47]. GST is well known to catalyse the conjugation of GSH to a range of substrates resulting in detoxification. GPx, together with GST, is vital in H_2_O_2_ metabolism. Thus, the decrease in the activity of GPx and GST, and a rise in the level of H_2_O_2_ observed in the present study in codeine-treated rabbits show that codeine led to peroxide-induced testicular damage. SOD is well known to catalyse the rapid dismutation of superoxide radicals to H_2_O_2_ and dioxygen while catalase converts the formed H_2_O_2_ into water and molecular oxygen. The significant reduction in the activities of testicular SOD and catalase observed in codeine-treated rabbits in the present study inarguably contributes to the excess H_2_O_2_ seen. Therefore, the overall decline in the glutathione system, SOD and catalase activities triggers robust oxidative stress.

There is growing evidence that oxidative damage permanently occurs to lipids of cellular membranes, proteins, and DNA. In nuclear and mitochondrial DNA, 8-OHdG is a predominant form of free radical-induced oxidative lesions and is a biomarker for oxidative DNA damage [48]. The rise in the level of this biomarker in codeine-treated animals reveals that codeine use promotes testicular oxidative DNA damage.

This study revealed that administration of codeine does not just trigger oxidative damage; it also up-regulates testicular nitric oxide (NO) and cysteine-aspartic proteases (caspase) 3. The overwhelming oxidant stress observed following codeine treatment ultimately excites an inflammatory reaction evident by a rise in testicular NO. It may be secondary to stimulation of NF-kB and enhancement of inducible nitric oxide synthase (iNOS) [49, 50], essential factors in NO production. NO couples with O_2_^-^ to form peroxynitrite (ONOO_2_^-^) which is hugely more reactive and harmful as compared to NO or O2^-^ alone. Nitric oxide has also been documented to trigger lipid peroxidation, protein degradation, and DNA damage [50-52]. Hence, the enhancement of testicular NO seen in codeine treatment also accounts for testicular damage.

Caspases are significant mediators of apoptosis. They are synthesized as pro-enzymes and activated by internal and external stimuli [53]. Caspase 3 is the most critical effector caspase, and the activation of its pathway is a hallmark of apoptosis used as a marker of death cascade [54, 55]. Findings from this study revealed that codeine up-regulated testicular caspase three expressions. The protease may cleaved inhibitor of caspase-activated DNAse I (ICAD) which after then stimulate caspase-activated DNAse I (CAD) [56, 57]. CAD is known to degrade chromosomal DNA within the nucleus and cause chromatin condensation, resulting in DNA fragmentation and cell apoptosis.

Activities of testicular enzymes ALP, ACP, LDH and γ-GT are markers of spermatogenesis [58]. They are indices of Sertoli function. The increased activities of ALP and ACP in codeine-administered animals may reflect the release of these non-specific phosphatases from the lysosomes of the degenerating cells and rapid catabolism of the testicular cells [59]. Codeine-induced increased activity of LDH observed in this study reflect testicular degeneration. It is likely due to a compensatory adaptation in an attempt to improve spermatogenesis and testicular regeneration from oxidative damage. γ-GT is a primary index of Sertoli function, and its activity is parallel to Sertoli cell maturation and replication [60]. The marked rise in γ-GT activity in codeine-treated animals in our present study suggests impaired Sertoli cell function and spermatogenesis, manifested by gross testicular atrophy, poor histoarchitecture of the testis, cessation of spermatogenesis, and germ cell maturation arrest.

The dose-dependent diminution in circulatory and intra-testicular concentrations of testosterone in codeine-treated rabbits is obviously due to oxidative damage and caspase-dependent apoptosis of the testes with associated testicular atrophy. Also, suppression of the levels of this androgen explains the rise in LH and FSH observed in codeine-treated animals, possibly via negative pituitary feedback. Spermatogenesis requires a high level of intra-testicular testosterone [61, 62]. In the present study, the cessation of spermatogenesis and the arrest of germ cell maturation seen in codeine-treated animals is a reflection of the reduced testicular level of testosterone. FSH has been documented to stimulate pyruvate and lactate production in Sertoli cells [63], while LDH catalyzes the interconversion of pyruvate and lactate together with the concomitant interconversion of NADH and NAD^+^. Thus, the raised FSH level in the current study in codeine-treated animals may account for the increased LDH activity seen.

T/LH and T/E_2_ are hormonal fertility indices. T/LH is a known index of Leydig cell function [64], while high T/E_2_ is a pointer to fertility [62]. Assessment of T/LH provides more meaningful information than comparisons of testosterone and LH levels separately [64]. In this study, we found that these hormonal fertility indices decreased in codeine-treated rabbits, indicating impaired Leydig cell function, which was manifested by suppression of both circulatory and intra-testicular levels of testosterone.

## Conclusion

Our current study reveals a novel mechanistic pathway for codeine-induced androgen suppression. Codeine depletes testicular glutathione, SOD, and catalase resulting in increased ROS production and inflammatory response which in turn leads to testicular lipid peroxidation, protein denaturation/inactivation, and DNA damage, with consequent testicular dysfunction and caspase-mediated testicular apoptosis.

## Declarations

### Abbreviations

Not applicable

### Ethics approval and consent to participate

The experiment followed the US NAS guidelines and ethical approval obtained from the Ethics Review Committee, Ministry of Health, Oyo State, Nigeria, with Reference Number AD13/479/1396.

### Consent for publication

All authors agree to the publication of this article

### Availability of data and material

Not applicable

### Authors’ contributions

ARE and AAF conceptualized and designed the study. ARE did the first draft. AAF reviewed the first draft.

## Acknowledgements

Not applicable

